# Tysserand - Fast and accurate reconstruction of spatial networks from bioimages

**DOI:** 10.1101/2020.11.16.385377

**Authors:** Alexis Coullomb, Vera Pancaldi

## Abstract

**Summary:** Networks provide a powerful framework to analyze spatial omics experiments. However, we lack tools that integrate several methods to easily reconstruct networks for further analyses with dedicated libraries. In addition, choosing the appropriate method and parameters can be challenging.

We propose *tysserand*, a Python library to reconstruct spatial networks from spatially resolved omics experiments. It is intended as a common tool to which the bioinformatics community can add new methods to reconstruct networks, choose appropriate parameters, clean resulting networks and pipe data to other libraries.

**Availability and implementation:** *tysserand* software and tutorials with a Jupyter notebook to reproduce the results are available at https://github.com/VeraPancaldiLab/tysserand

**Supplementary information:** Supplementary data are available at *Bioarxiv* online.

## 1 Introduction

Recent technologies have made it possible to produce phenotypic data at the resolution of single cells (or higher) in intact sample slices, both at the levels of proteins [1, 2] or mRNA [3, 4]. Taking advantage of spatial information is essential for revealing the biology of healthy organs and dissecting the complex processes involved in cancer, such a tumor progression and response to treatments[5].

Existing spatial omics analysis libraries such as trendsceek[6], SpatialDE[7] and PySpacell[8] use marked point processes theory. Another fruitful approach is to represent tissues as networks, where nodes are cells and edges are interactions between cells which are established through physical contact. Network theory is already used for spatial analysis in the Python Spatial Analysis Library (PySAL)[9] for geospatial data science, and PySpacell, based on PySAL, provides 3 methods to reconstruct networks: k-nearest neighbors, radial distance neighbors and cell contact neighbors. However, due to its dependence on PySAL, it is not ideally suited to test other network reconstruction methods and PySAL methods do not scale well with big datasets of thousands of cells. Moreover, the choice of a reconstruction method and parameters can be hard, and the potential to use the reconstructed network with other dedicated network analysis libraries remains a priority. Here, following the Unix philosophy[10] according to which *programs do one thing and do it well* and *programs work together*, we present *tysserand*, a Python library to reconstruct spatial networks starting from object positions (cells, nuclei, …) or image segmentation results. We aim at encouraging the centralization of efforts in network construction from the bioinformatics community in one place, to promote integration of new reconstruction methods, providing algorithms for parameters selection, network processing to remove reconstruction artifacts, computational performance improvement and the addition of interfaces with external network analysis libraries such as NetworkX[11], iGraph[12] or Scanpy[13] for single-cell data analysis.

## 2 Results

*tysserand* can consider two types of alternative inputs (Figure 1). First, an M×2 array of the M cells’s x/y coordinates or, second, a ‘segmentation image’ with integers ranging from 0 to K representing K segmented areas in a microscopy image, with 0 values indicating the background. For the coordinates array input, 3 methods are already available to reconstruct networks, based on the Scipy[14] library implementation: k-nearest neighbors (knn), radial distance neighbors (rdn) and Delaunay triangulation (Figure S3). We think Delaunay triangulation is best suited to represent tissues and interactions between contacting cells, whereas the rdn method is more appropriate to model interactions by diffusing chemicals, as already noticed by PySpacell authors[8]. For inputs consisting in cell segmentation images, we implemented the area contact neighbors method (Figure S7). It leverages the scikit-image[15] and OpenCV[16] libraries to detect for each area which neighbors are in direct contact or closer than a given distance. The *tysserand* library provides simple visualization utilities to choose appropriate parameters for each network construction method and tools to clean the resulting networks from typical artifacts, such as very long edges between nodes on the border of samples after Delaunay triangulation (Figure S4). *tysserand* also provides utilities to visualize interactively multi-channels bio-images with the Napari[17] library, which allows the user to pan, zoom and modify each channel intensity to better define or modify networks built with *tysserand* (Figure S2).

**Figure 1:**
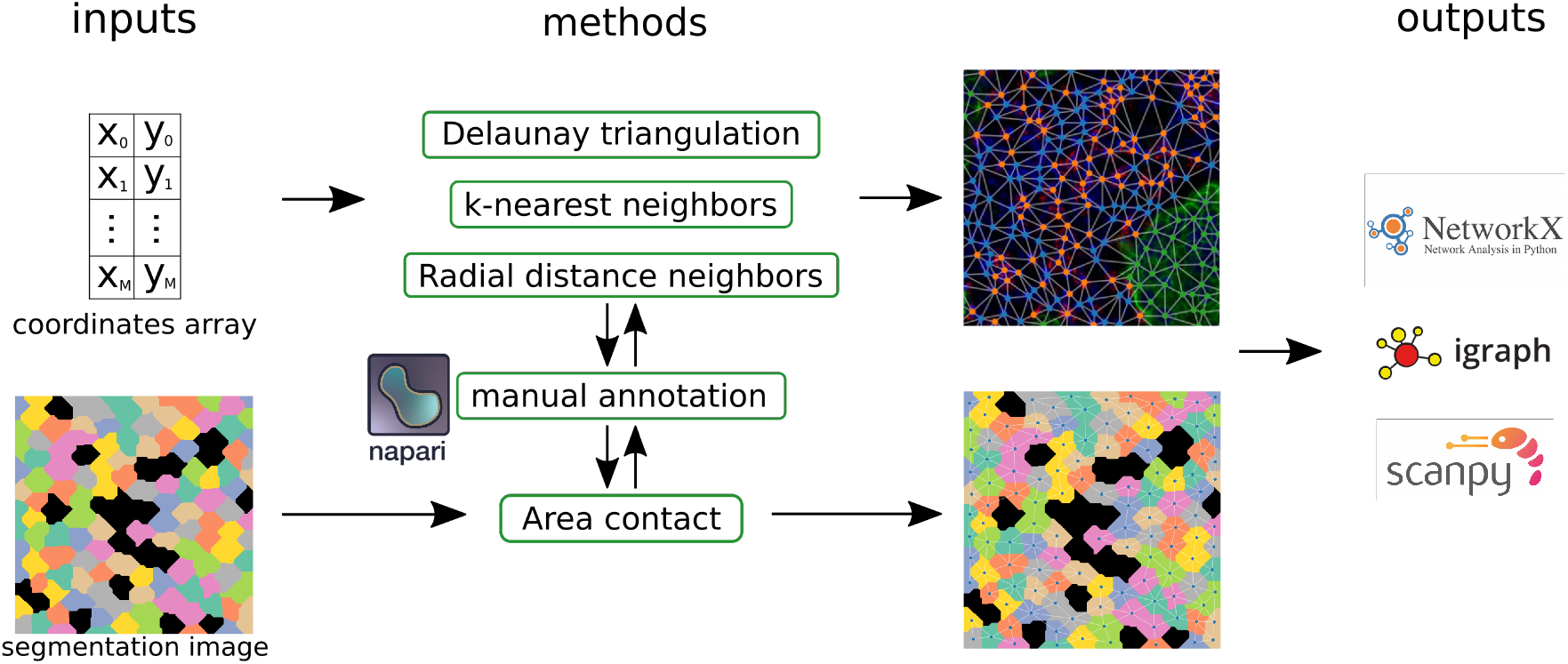
*tysserand* can take as inputs an array of nodes positions or an image resulting from segmentation processing. 4 methods are implemented for now to reconstruct spatial networks, and they can be manually curated with napari for interactive visualization and annotations. The resulting networks can then be exported to formats compatible with libraries dedicated to network analysis or using networks in their downstream analyses.

Internally, *tysserand* adopts simple and efficient representations of networks to allow rapid prototyping of new network construction or cleaning methods. A network is represented by 2 arrays: a first M×2 array for nodes coordinates (that are the center of segmented objects if a segmentation image is provided as input), and an L×2 array to represent the L edges between nodes indicated by their index in the first coordinates array. Finally, *tysserand* can convert networks into formats used by specialized libraries such as NetworkX, iGraph and Scanpy.

To compare the quality of reconstructed networks across methods, we considered the network reconstruction task as a classification task, were a model has to predict the presence or absence of edges between all possible pairwise nodes combinations. We can then define the true positives as the edges that were successfully predicted to exist, the false positive as the edges that were erroneously predicted to exist, the false negative as the edges that were erroneously predicted to not exist, and the true negatives as the edges that were successfully predicted to not exist. We will note their quantity TP, FP, FN and TN respectively. Given n the number of nodes in a network, TN scales with the number of all pairwise combinations, which is 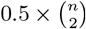, whereas the number of actual edges scales linearly with the number of nodes, as cells have a limited number of possible neighbors, regardless the size of the tissue sample. Thus, as the size of the network increases, TP, FP and FN become negligible compared to TN, which prevents us from using standard classification quality metrics such as the adjusted accuracy or the Matthews correlation coefficient. Thus, we defined a measure of network reconstruction quality as the *true positive ratio* 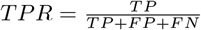.

*tysserand* allowed us to easily compare networks generated with the knn and rdn methods, the Delaunay triangulation, as well as the equivalent Voronoi tessellation implemented in PySAL.

We first compared the execution time of these methods on randomly generated sets of node positions (Figures S6). Across all sizes of node sets the *tysserand* Delaunay implementation is always at least 51 times faster than PySAL’s. This speed-up is valuable in the range of several tens of thousands of nodes, which is often the size of current bioimage datasets.

We then compared the quality of reconstructed networks on simulated tissue images (data available on the GitHub repository, description in Text S1, Figures S7 and S8). For the most realistic simulations implementing noise in cell positions and empty spaces in the tissue, the default *tysserand* Delaunay method is always more accurate than PySAL method (Table S1). For less realistic simulations performances are discussed in Text S1. Finally, we compared the methods’ output quality on a real bioimage (Figure S1) annotated with napari and the associated utilities implemented in *tysserand*. Qualitatively, Delaunay triangulation is best suited to reconstruct spatial networks of tissue samples, as most edges link contacting cells, whereas the knn and rdn methods produce networks with excessively connected areas or missing edges (Figure S3). Quantitatively, the *tysserand* Delaunay method was the most accurate method (Table S2), slightly better than PySAL’s method while being 126 times faster.

## 3 Conclusion

*tysserand* can reconstruct spatial networks from different inputs, such as sets of node positions or segmented areas, and the resulting networks can be further processed with dedicated network analysis libraries. It already implements 4 common network reconstruction methods as well as tools to facilitate the choice of parameters for network construction, artifact removal and tools to facilitate manual networks creation or modifications. *tysserand* Delaunay triangulation is faster than the PySAL equivalent method even with big datasets, which is important for the analysis of multiple tissue samples, and produces networks of better quality on real bioimages. We hope the bioinformatics community will be willing to participate in the implementation of new methods for more accurate and domain specific spatial network reconstruction to advance the field of bioimage processing.

## Acknowledgements

We thank Professor Pierre Brousset and the Imag’IN facility at IUCT Oncopole, Toulouse for providing the multiplex immuno-fluorescence images. We also thank the referee whose suggestions significantly contributed to improve this manuscript.

