## Supplementary Material for "Tysserand - Fast and accurate reconstruction of spatial networks from bioimages"

Alexis Coullomb and Vera Pancaldi
Centre de Recherches en Cancérologie de Toulouse, Toulouse, France

Text S1: We compared the execution time of the tysserand Delaunay triangulation and the mathematically equivalent Voronoi tessellation implemented in PySAL on randomly generated sets of node positions. The PySAL implementation, being based on shape objects, does not produce long edges artifacts (Figure S5). Since the tysserand Delaunay method does produce this type of artifacts, we also benchmarked tysserand's method including automated edge trimming, which could slow down the performance but is necessary to obtain results that are comparable to PySAL. Across all sizes of node sets, the tysserand implementation of the Delaunay triangulation, including the automated network artifact removal option, is always at least 51 times faster than PySAL's (Figure S6). It is likely that the lower performance of the PySAL library is due to the use of 'shape' objects, which are common and important in geographical sciences but compromise the algorithm scalability. Since biological image applications often do not require the concept of 'shape', considerable improvements in computational speed can be obtained using tysserand.

Since the aim of tysserand is to work on real biological images, we adopted an approach that starting from a regular lattice can add realistic features to this benchmark image, representing the non-perfect shape of cells and tissue topological arrangements characteristic of real biological samples.

We explored two parameters that can be varied to simulate more realistic images: `hole_proba` that is the probability to delete one segmentation object to mimic natural empty spaces in tissues, and `noise_sigma` that controls the level of gaussian noise added to initial objects position before defining their segmentation area.

On simulated data, the KNN method is overall the worst performing one, whereas the RDN is overall the best performing method with the optimal radius parameter given from a priori knowledge (Table S1). On ideal regular lattices (top 6 rows, no noise no empty space) PySAL equals the RDN method followed closely by Delaunay method with the 3 possible trimming options. With the most realistic simulated images, tysserand's Delaunay triangulation with adaptive edge trimming is the best performing method after the RDN method, but Delaunay method doesn't require a priori knowledge to select an optimal parameter, which is essential for real applications in which the ground truth is not known.

When comparing the quality of reconstructed networks for tysserand methods and PySAL Voronoi method on the real multiplex immuno-fluorescence bioimage, the tysserand Delaunay triangulation with adaptive edge trimming is the best performing method, followed by PySAL's method (Table S2). On this image this tysserand method was 126 times faster than PySAL's method.

We qualitatively assessed the differences between the resulting networks of the Delaunay triangulation, knn and rdn methods by visual inspection, and we observed the following (Figure S3):

- The Delaunay triangulation with edge trimming produces a network that looks similar to what we can expect from a tissue, i.e., most edges link contacting cells, as visible on the mIF image (Figure S3 b)
- rdn produces excessively connected areas where the density of nodes is high
- knn produces a network with missing edges where we could expect them based on cell contacts, as well as edges passing through neighboring cells, which is not suitable to model interactions dependent on direct physical contact.

We thus conclude that Delaunay triangulation is the most suitable method for these biological tissue images.

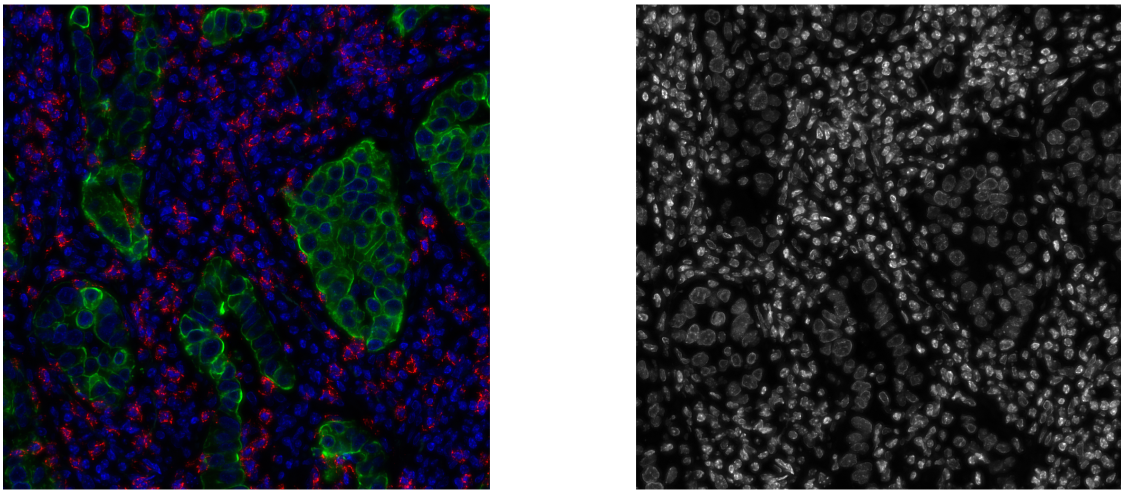

(a) (b)

Figure S1: Exemplary tissue image (a) extracted from a multiplex immunofluorescence (mIF) *Whole Slide Image* of a lung tumor biopsy. Pseudo colors are blue: DNA, green: PanCK (tumor), red: CD8 (cytotoxic T cells). Nodes positions are defined by manual marking of nuclei (shown in b), nodes class (blue / green / red color coded in the following document) are defined from markers in image (a).

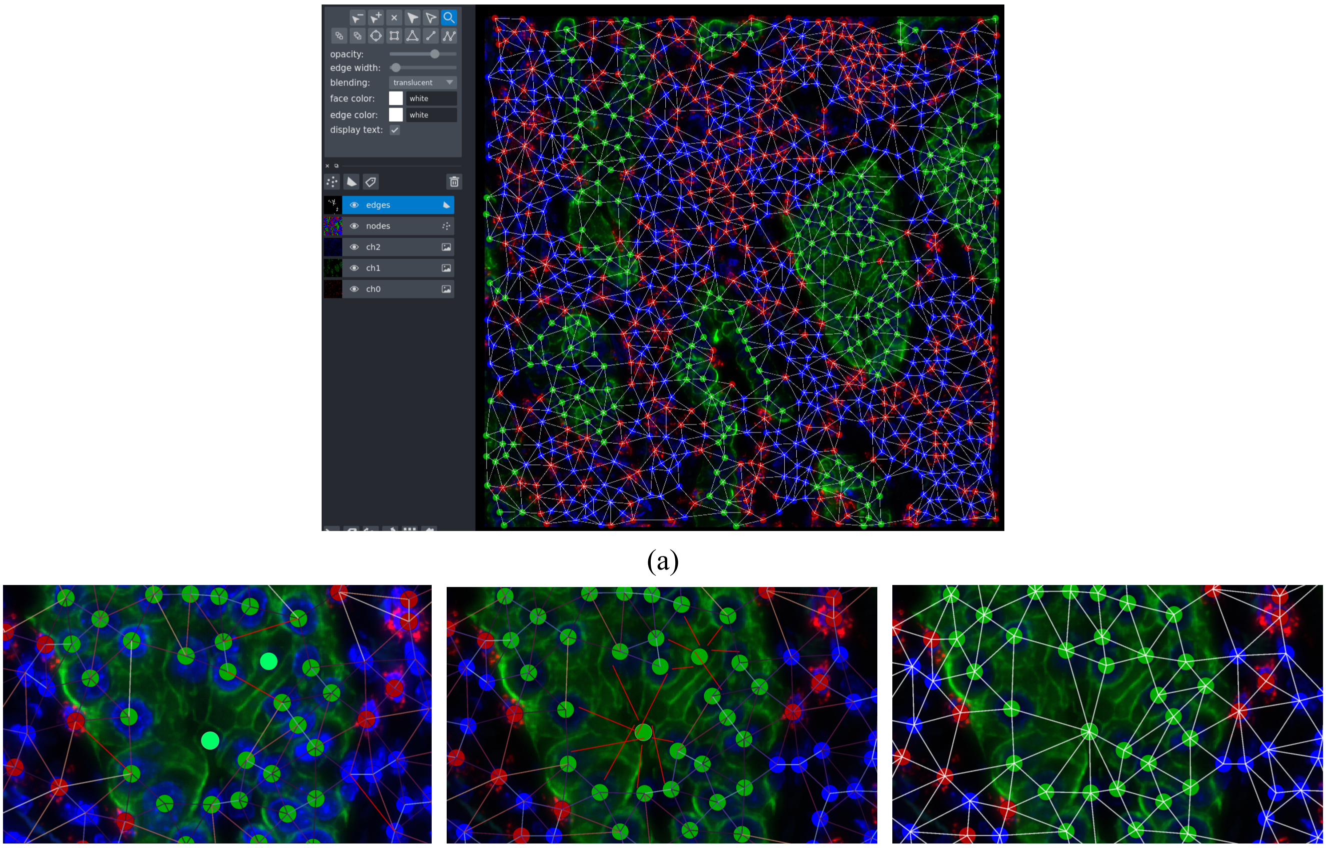


(a)

(b) (c) (d)

Figure S2: tysserand includes tools to facilitate interactive network visualisation and annotation with napari (a). Nodes and edges can be added and removed manually and saved for later analysis. In this example all nodes were added manually to generate a benchmark dataset, then the tysserand Delaunay method was used to make a first set of edges. (b) After increasing the DAPI channel we can detect missing nodes that we add (light green) and edges that should be discarded (red). (c) We can then manually add new edges (red). (d) tysserand includes a method to assign each edge to its pair of nodes in order to build the network in the tysserand format, and to clean the position of edges that were manually added.

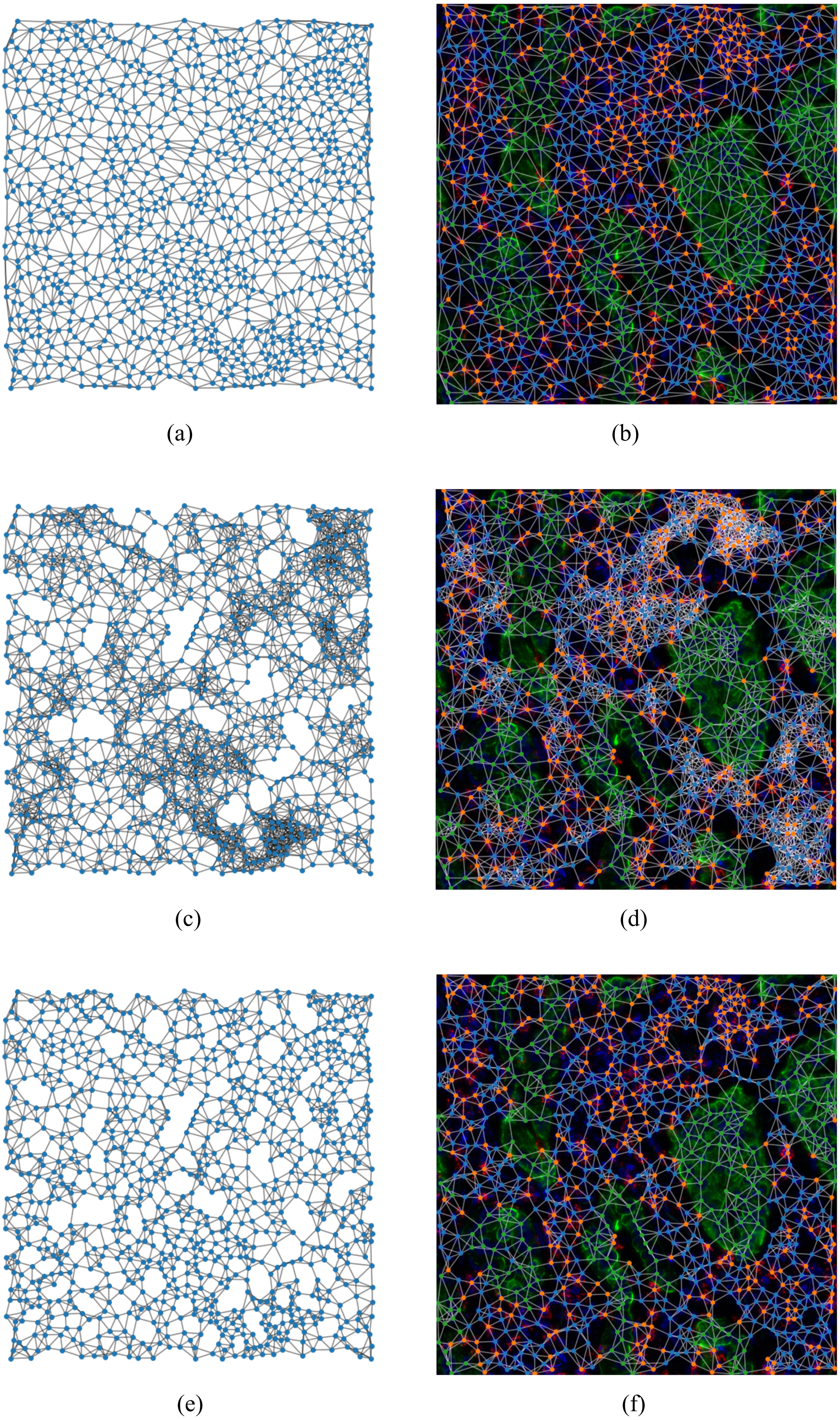


Figure S3: Delaunay triangulation after edge trimming (a) and resulting network superimposed on the mIF image (b). Radial distance neighbors (r=60) network (c) and network superimposed on the mIF image (d). The variability in cell density induces very lowly and very highly connected areas. k-nearest neighbors (k=6) network (e) and network superimposed on the mIF image (f). Some areas are less connected than expected given the apparent cells contacts, and some edges go *through* neighbors.

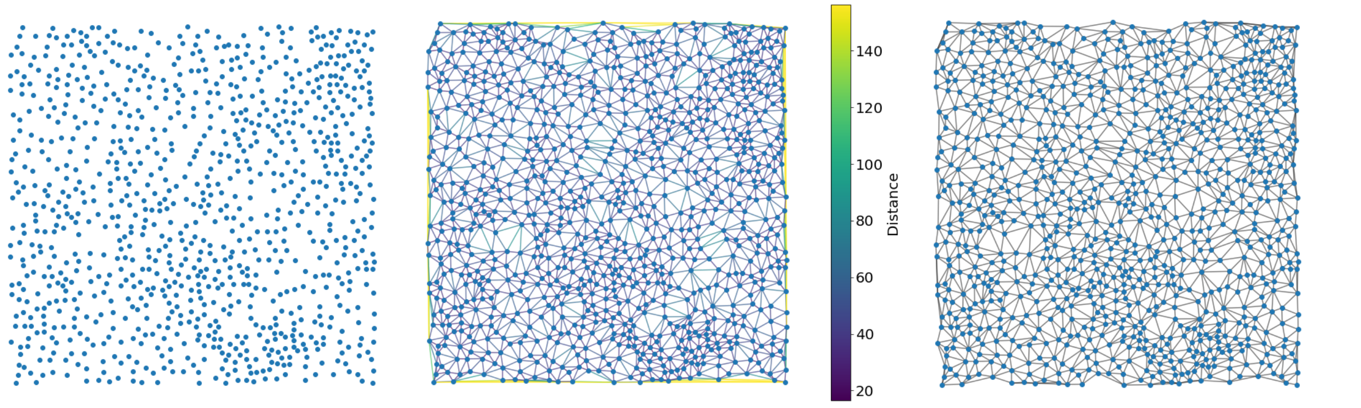


(a) (b) (c)

Figure S4: (a) Scatter plot of 385 nodes, their coordinates are given by a 385x2 array. (b) Delaunay triangulation can produce long edges artifacts, most often at the border of the samples (yellow edges). Users can trim edges longer than a manually defined distance using distance plots like in (b), or define this threshold at a distance corresponding to the 99th (or other) percentile of lengths., or choose the default method that adapts this percentile threshold given the number of nodes in the network (c) The resulting cleaned network after edge trimming.

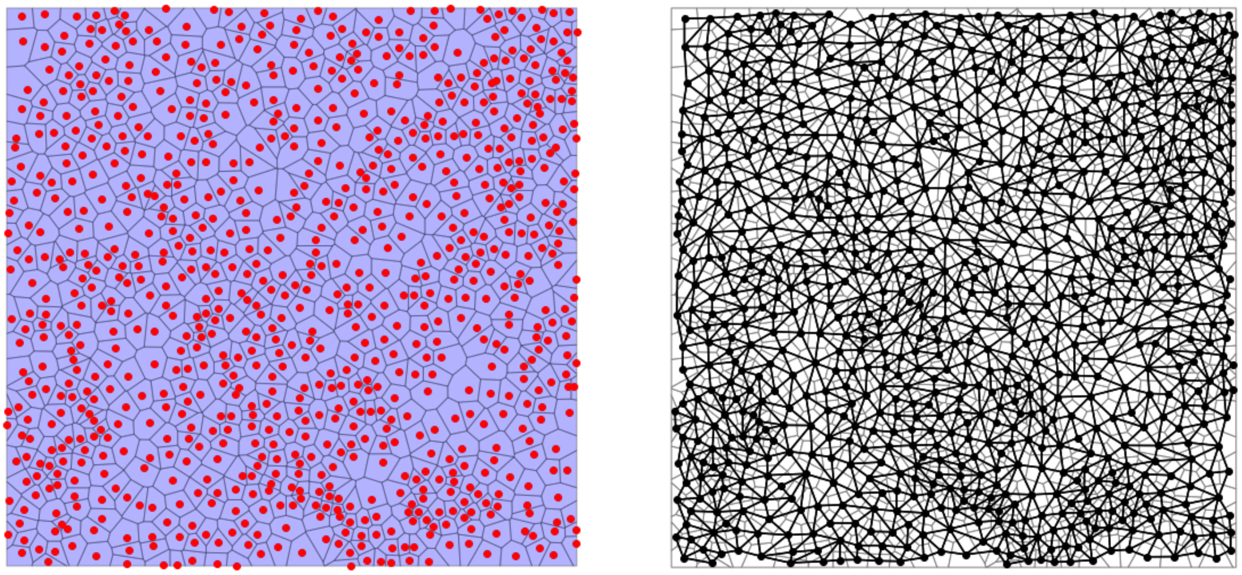

(a) (b)

Figure S5: (a) PySAL implementation of Voronoi tessellation results in 'shape objects' that are common in geosciences. (b) Contacts are computed between these objects and they define the Voronoi diagram (equivalent to Delaunay triangulation). Thanks to the use of contacts between shape objects, there are no long edges artifacts.

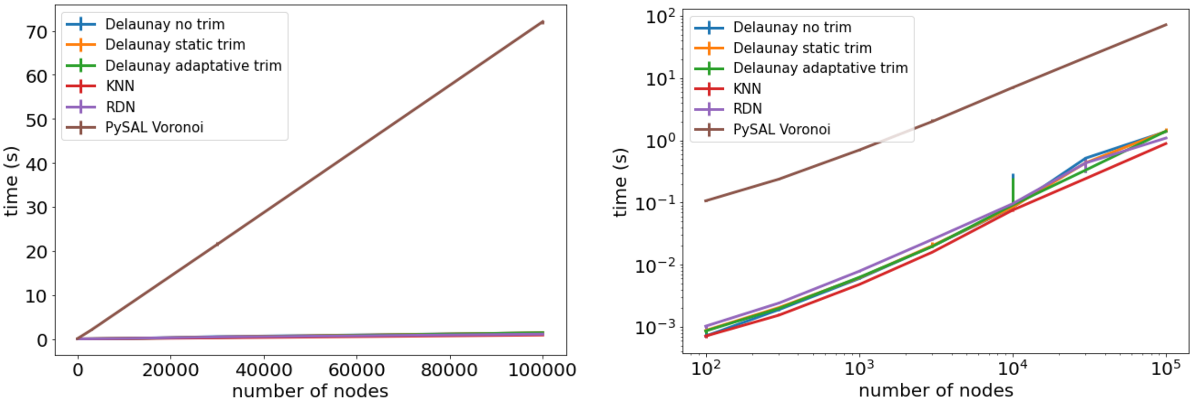


(a) (b)

| size | 100 | 300 | 1000 | 3000 | 10000 | 30000 | 100000 |
|---|---|---|---|---|---|---|---|
| **tysserand** | 0.000863 | 0.00199 | 0.00626 | 0.0197 | 0.0909 | 0.333 | 1.4 |
| **PySAL** | 0.107 | 0.238 | 0.698 | 2.01 | 7.04 | 21.4 | 72.1 |
| **speed up** | 124 | 120 | 111 | 102 | 77.5 | 64.4 | 51.4 |

(c)

Figure S6: Execution time comparison between Delaunay/Voronoi implementations of tysserand (based on scipy) and PySAL for various sizes of sets of nodes. Tested sizes are 100, 300, 1000, 3000, 10000, 30000 and 100000 nodes. For each size, 10 random sets are generated and both implementations are tested on the same sets. Graphs display on uniform scale (a) or log-log scale (b) the median of execution time (s), error bars are the first and third quartiles. (c) Table of benchmark results, tysserand is at least 51 times faster than PySAL for Delaunay triangulation.

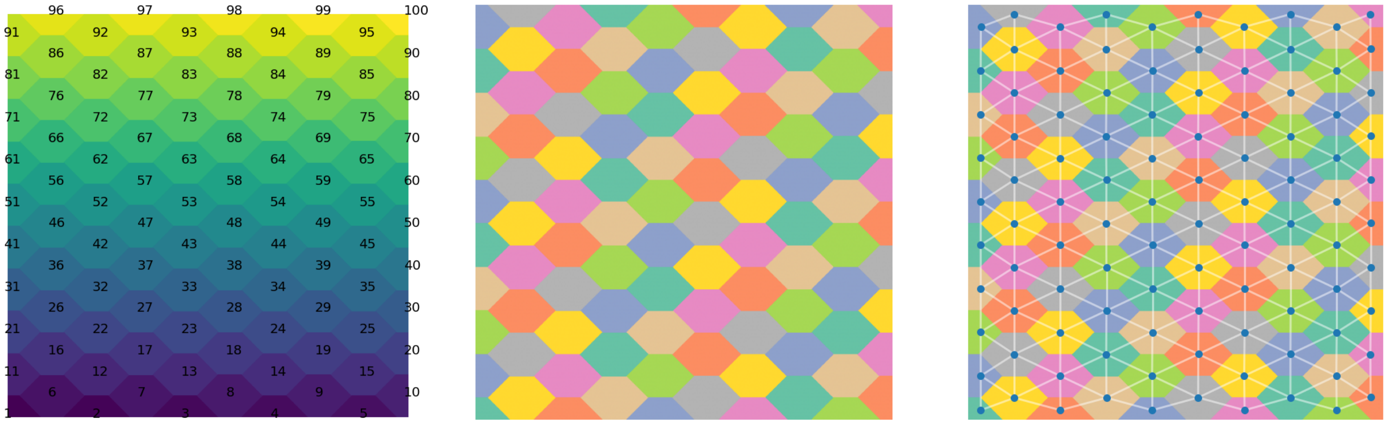

(a) (b) (c)

Figure S7: A segmentation image (a) is a 2D array containing areas of integers from 0 to K resulting from the detection of K areas (cells, nuclei, …). Here, areas are color-coded, and their integer value is displayed as the contrast between adjacent areas can be very low. We use a scikit-image function to display areas with a better contrast (b). The area contact method uses scikit-images and OpenCV functions to detect for each area the areas that are in direct contact or closer than a user-defined distance (c). On this generated set of tiles mimicking a tissue, the area contact method appears well suited to reconstruct tissue networks.

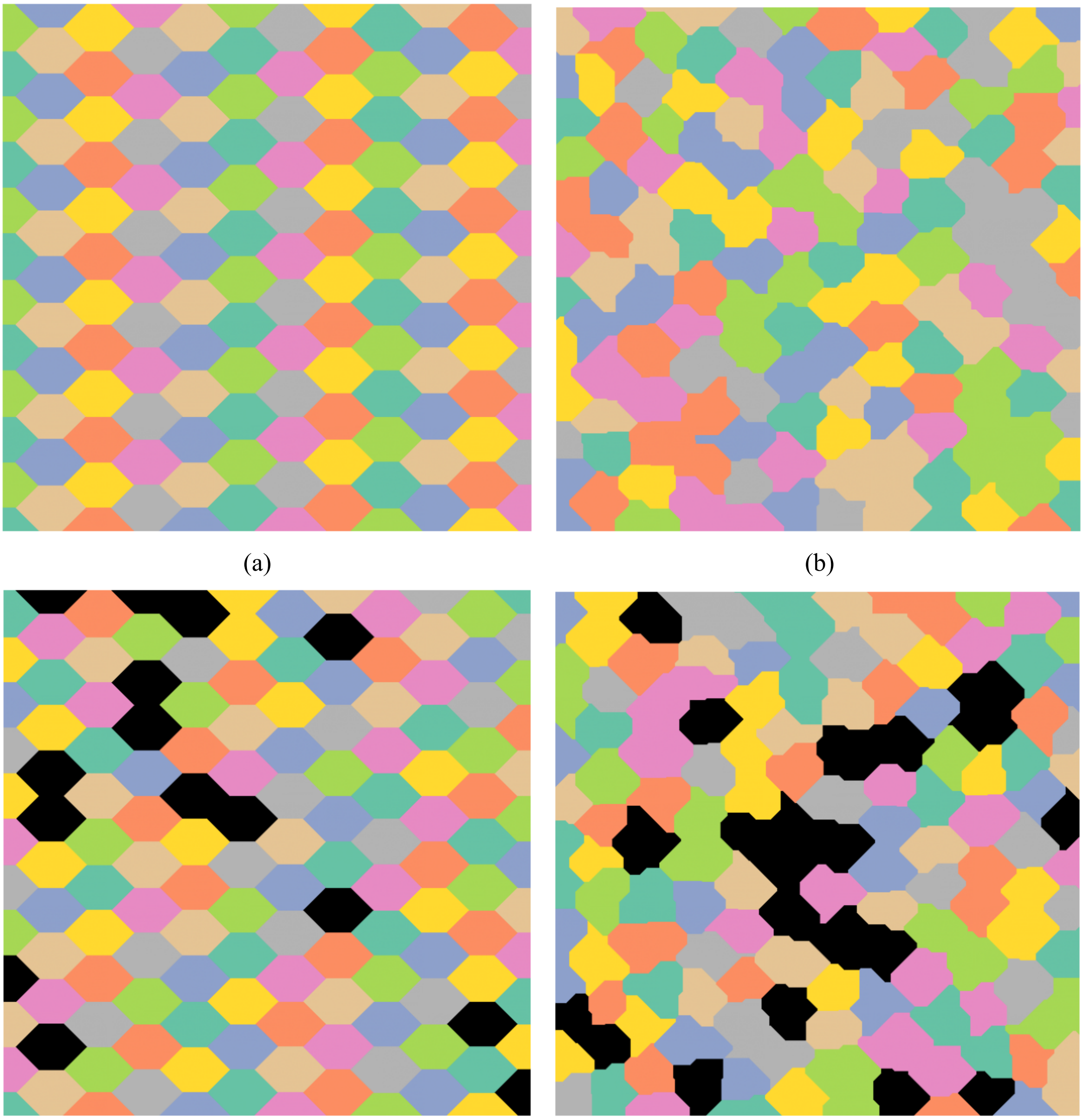

(a) (b) (c) (d)

Figure S8: Tissue images generated with increasing realism. Left and right columns: images simulated with noise_sigma = 0 or 10 respectively. Top and bottom rows: images simulated with hole_proba = 0 or 0.1 respectively.

| hole_proba | noise_sigma | size | TP ratio Delaunay no trim | TP ratio Delaunay static trim | TP ratio Delaunay adaptative trim | TP ratio KNN | TP ratio RDN | TP ratio PySAL Voronoi |
|---|---|---|---|---|---|---|---|---|
| 0.0 | 0.0 | 500 | 0.929 | 0.939 | 0.843 | 0.811 | 1.000 | 1.000 |
| | | 867 | 0.961 | 0.984 | 0.844 | 0.860 | 0.996 | 1.000 |
| | | 1582 | 0.970 | 0.980 | 0.950 | 0.932 | 0.999 | 1.000 |
| | | 2739 | 0.984 | 0.996 | 0.971 | 0.901 | 1.000 | 1.000 |
| | | 5000 | 0.988 | 0.998 | 0.990 | 0.892 | 1.000 | 1.000 |
| | | 8661 | 0.994 | 0.995 | 0.995 | 0.965 | 1.000 | 1.000 |
| | 10.0 | 500 | 0.897 | 0.907 | 0.868 | 0.803 | 0.946 | 0.974 |
| | | 867 | 0.905 | 0.915 | 0.922 | 0.830 | 0.921 | 0.958 |
| | | 1582 | 0.925 | 0.934 | 0.944 | 0.850 | 0.935 | 0.955 |
| | | 2739 | 0.937 | 0.946 | 0.950 | 0.861 | 0.939 | 0.956 |
| | | 5000 | 0.940 | 0.949 | 0.949 | 0.864 | 0.931 | 0.951 |
| | | 8661 | 0.944 | 0.950 | 0.950 | 0.867 | 0.930 | 0.950 |
| 0.1 | 0.0 | 500 | 0.855 | 0.865 | 0.917 | 0.787 | 1.000 | 0.928 |
| | | 867 | 0.868 | 0.878 | 0.982 | 0.851 | 0.995 | 0.910 |
| | | 1582 | 0.886 | 0.895 | 0.950 | 0.897 | 0.999 | 0.918 |
| | | 2739 | 0.884 | 0.894 | 0.920 | 0.896 | 1.000 | 0.902 |
| | | 5000 | 0.896 | 0.909 | 0.916 | 0.903 | 1.000 | 0.908 |
| | | 8661 | 0.925 | 0.956 | 0.956 | 0.935 | 1.000 | 0.931 |
| | 10.0 | 500 | 0.822 | 0.832 | 0.929 | 0.780 | 0.949 | 0.894 |
| | | 867 | 0.823 | 0.832 | 0.928 | 0.816 | 0.923 | 0.873 |
| | | 1582 | 0.838 | 0.847 | 0.896 | 0.841 | 0.930 | 0.869 |
| | | 2739 | 0.847 | 0.856 | 0.881 | 0.855 | 0.937 | 0.866 |
| | | 5000 | 0.857 | 0.865 | 0.875 | 0.866 | 0.931 | 0.867 |
| | | 8661 | 0.858 | 0.867 | 0.868 | 0.869 | 0.928 | 0.864 |

Table S1

Quality of reconstructed networks for tysserand methods and PySAL Voronoi method for increasing size and realism.

| | Delaunay no trim | Delaunay static trim | Delaunay adaptative trim | KNN | RDN | PySAL Voronoi |
|---|---|---|---|---|---|---|
| TP ratio | 0.931 | 0.941 | 0.977 | 0.749 | 0.648 | 0.954 |
| time (s) | 0.00951 | 0.00752 | 0.00711 | 0.00507 | 0.00847 | 0.898 |

Table S2

Quality of the reconstructed network of the mIF bioimage and execution time for tysserand methods and PySAL Voronoi method.

Text S2: Pseudo code of reconstruction methods.

```
FUNCTION build_delaunay:
      pairs = CALL scipy voronoi and extract coordinates
      IF we trim edges:
      distances = CALL distance_neighbors with coordinates and pairs
      IF trim parameter is not a distance but a method:
             trim parameter = CALL find_trim_dist with distances and method parameters
      pairs =  all pairs which edge length is below trim_parameter
      ENDIF
      ENDIF
      RETURN pairs

FUNCTION distance_neighbors:
      coordinates of first point = select coordinates of the first node for all pairs of nodes
      coordinates of second point = select coordinates of the second node for all pairs of nodes
      distances = compute euclidean distance between first points and second points
      return distances

FUNCTION find_trim_dist:
      IF method is 'percentile_size':
      proportion of border edges = 4 / SQRT(number of nodes)
      percentile value = 100 * (1 - prop_edges * 0.5)
      distance threshold = CALL numpy percentile with distances and percentile value
      ELSE IF method is 'percentile':
      distance threshold = CALL numpy percentile with distances and percentile value
      ENDIF
      RETURN distance threshold

FUNCTION build_knn:
      tree = CALL scipy BallTree with coordinates of nodes
      indices = CALL query method of tree to make a matrix of size number of nodes x number of
neighbors
      pairs = CALL pairs_from_knn with indices to make a matrix of shape number of pairs x 2
(first and second node ids)
      RETURN pairs

FUNCTION pairs_from_knn:
      number of pairs = comp
      source_nodes = array of nodes indices repeated k time (number of neighbors)
      target_nodes = select neighbors of each node and make an array of them
      pairs = merge source_nodes and target_nodes
      sort in increasing order nodes ids for each pair of nodes
      drop duplicate pairs
      RETURN pairs


FUNCTION build_rdn:
      tree = CALL scipy BallTree with coordinates of nodes
      indices = CALL query method of tree to make a list of size number of nodes containing
arrays with neighbors indices
      # clean arrays of neighbors from self referencing neighbors
      # and aggregate at the same time
      source_nodes = empty list
      target_nodes = empty list
      FOR source_node_id, array_neighbors in indices:
      neighbors = array_neighbors without source_node_id
      ADD to source_nodes a list of repeated source_node_id of length the number of neighbors
      ADD to target_nodes the list of neighbors
      ENDFOR
      flatten arrays of arrays for source_nodes
```

```
      flatten arrays of arrays for target_nodes
      pairs = merge source_nodes and target_nodes
      sort in increasing order nodes ids for each pair of nodes
      drop duplicate pairs
      RETURN pairs


FUNCTION build_contacting:
      source_nodes = empty_list
      target_nodes = empty_list
      FOR mask_id in list of segmentation object ids:
      neighbors = CALL find_neighbors with segmentation image, mask_id, radius
      ADD to source_nodes a list of repeated source_node_id of length the number of neighbors
      ADD to target_nodes the list of neighbors
      ENDFOR
      flatten arrays of arrays for source_nodes
      flatten arrays of arrays for target_nodes
      pairs = merge source_nodes and target_nodes
      sort in increasing order nodes ids for each pair of nodes
      drop duplicate pairs
      RETURN pairs

FUNCTION find_neighbors:
      mask = make binary image with pixels belonging to the object are set to 1, 0 otherwise
      # create the border in which we'll look at other masks
      kernel = create binary kernel according to the specified radius
      dilated = CAL openCV dilate with mask and kernel
      dilated = convert dilated to boolean
      neighbors = select unique id values in masks where dilated is True
      neighbors = neighbors without the initial cell id of interest
      neighbors = neighbors without background value
      RETURN neighbors

FUNCTION mask_val_coord:
      coordinates = CALL scikit-image measure.regionprops_table with masks to get centroids
coordinates and masks ids
      coordinates = organize coordinates as a DataFrame
      return coordinates

FUNCTION refactor_coords_pairs:
      mapper = dictionary to relate mask ids to indices of nodes in the coordinates DataFrame
      pairs = DataFrame with source and target nodes for each edge
      source nodes in pairs = convert source nodes in pairs from masks ids to nodes ids
      target nodes in pairs = convert target nodes in pairs from masks ids to nodes ids
      coordinates = extract array of the coordinates DataFrame
      pairs = extract array of the pairs DataFrame
      RETURN coordinates, pairs
```
